# Walking entrains unique oscillations for central and peripheral visual detection

**DOI:** 10.1101/2024.07.04.602020

**Authors:** Cameron K. Phan, Matthew J. Davidson, David Alais

**Affiliations:** School of Psychology, The University of Sydney

**Author notes:** This research is supported by an Australian Government Research Training Program (RTP) Scholarship to CP and an Australian Research Council grant (DP210101691) to DA.

**Keywords:** active perception, walking, locomotion, virtual reality, visual oscillations

## Abstract

It is important to investigate perception in the context of natural behaviour in order to reach a holistic account of how sensory processes are coordinated with actions. In particular, the effect of walking upon perceptual and cognitive functions has recently been investigated in the context of how common voluntary actions may dynamically impact upon visual detection. This work has revealed that walking can enhance peripheral visual processing, and that during walking, performance on a visual detection task oscillates through good and bad periods within the phases of the stride-cycle. Here, we extend this work by examining whether oscillations in visual detection performance are uniform across the visual field while walking. Participants monitored parafoveal (∼3.7 d.v.a) and peripheral (∼7 d.v.a) locations left/right of fixation for the onset of targets while walking at a natural pace in wireless virtual reality. For targets at all locations accuracy, reaction times and response likelihood oscillated within each individual’s stride-cycle, at primarily 2 or 4 cycles per stride. Importantly, oscillations in accuracy and reaction time shared the same frequency at both locations but were decreased in amplitude and phase-lagged in the periphery, revealing an interaction between visual field locations and oscillations in performance. Together, these results demonstrate that oscillations in visual performance entrained by the stride-cycle occur with unique amplitudes and phases across the visual field.

---

Perceptual research has traditionally required participants to complete tasks in highly controlled and stimulus-limited environments with restricted physical autonomy (e.g., with seated observers, and head stabilised on a chin rest). However, perception in daily life is typically far more dynamic with observers voluntarily self-generating actions (e.g., saccades, reaching, locomotion) which in turn alter the sensory inputs driving perceptual experience. This difference between experimental and real-life conditions reduces the ecological validity of previous work and has left the field of active perception in the context of everyday actions relatively unexplored.

The importance of every day actions in perception has been appreciated for many years (Bardy et al., 1996; Chapman et al., 1987; Gibson, 1955, 1962; Warren & Hannon, 1988) but technical challenges have been an impediment to empirically investigating natural behaviour. Recent advances in portable, wearable devices and virtual reality technology have removed many of these barriers, and active perception research can now be more easily conducted. For example, a number of recent studies involving observers performing perceptual tasks during walking have provided insight into the interaction between everyday actions and perception (Davidson et al., 2023, 2024; Matthis et al., 2017, 2018; Matthis & Fajen, 2014; Selinger et al., 2015).

Walking is a movement generated by most able-bodied individuals on a daily basis, facilitating locomotion through the environment to intended destinations (Winter, 1983). Walking is a very cyclic behaviour with well-established phases that are consistently observed and easily modelled across multiple performers (MacDougall & Moore, 2005). Walking is described in terms of steps, and phases of the stride-cycle (Kharb et al., 2011). A step begins with the lift-off of a foot (e.g., left) and ends with its return to the ground and during this period the other foot (e.g. right) supports the individual’s mass. A stride consists of two successive steps, beginning with the lifting of one foot and ending with the lifting of the same foot. The stride-cycle is often divided into two phases known as ‘stance’ and ‘swing’ phase. The stance phase begins following the heel-strike of one foot until the subsequent toe-lift of the same foot and includes a double-support phase while both feet are on the ground. The swing phase includes the stages when one foot is not in contact with the floor. The consistency of gait during walking has been widely studied and step rates across the population on smooth, level ground are clustered around 2 Hz (Gard et al., 2004; Hausdorff et al., 1996; Hirasaki et al., 1999; Moore et al., 2001; Pozzo et al., 1990). Within individuals, step rates are also highly consistent and rhythmic (MacNeilage, 2020; MacNeilage & Glasauer, 2017). Because walking is such a common daily action, is well studied biomechanically and involves consistent repetitions of a cyclical voluntary action, it is one of the most suitable movements for investigating the effects of action on perception.

When walking through an environment, an individual must remain alert for potential obstacles lying on the path ahead as well as monitor for risks and collisions peripheral to the intended course (Drew & Marigold, 2015; Matthis et al., 2017). With that in mind, the present study examines detection of visual targets in the central versus peripheral field in walking observers. Previous work by Cao and Händel (2019) found neurophysiological and behavioural results suggesting enhanced processing of peripheral visual input while walking (relative to stationary) using a contrast sensitivity task. Enhanced peripheral processing was implied by greater suppression of central visual targets from competing peripheral targets. This interesting work prompted our current study design which involves simple detection of brief targets in either a central or peripheral location which is more akin to detecting targets that may appear suddenly on a path or adjacent to it. We also introduce a new analysis in which detection performance is binned relative to the stages of the stride-cycle, so that any modulations of visual detection during walking can be revealed.

Previous experiments from our group found oscillations of visual detection performance when binning performance relative to the phases of the stride-cycle. These oscillations primarily occurred at low frequencies, such as 2 or 4 cycles per stride (*cps;* Davidson et al., 2024), that were entrained to the rhythm of individual footfall. Notably, these oscillations were recorded during a visual detection task that required fixating on a slowly drifting target region, precluding an analysis of stride-cycle effects based on target eccentricity. Previous work has also shown that rhythms in perception can propagate across retinotopic visual space (Sokoliuk & vanRullen, 2016; Fakche & Dugue, 2023), possibly supported by the spatial propagation of neural oscillations across the cortex (Lozano-Soldevilla & VanRullen, 2019; Prechtl et al., 2000). Here we investigated whether locomotion induced perceptual oscillations also propagate across retinotopic space, by comparing performance at central and peripheral target locations when binned according to the phases of an individual’s stride-cycle.

The current study employed a simple visual detection task during free walking using a wireless virtual reality (VR) headset. By using VR, we were able to simulate an outdoor setting that could be replicated across participants (Fink et al., 2007), resulting in a naturalistic optic flow that was absent when treadmills (e.g., Benjamin et al., 2018; Gramann et al., 2010) or stationary bicycles (e.g., Bullock et al., 2017) have been used in past research. The use of wireless VR also enabled the tight experimental control of visual stimuli, simultaneous recording of eye-movements, and the capture of changes in head height from which the phases of the stride-cycle were extracted (Davidson et al., 2024). Participants made speeded responses to targets that were presented at randomly chosen time points at either central or peripheral locations while they walked along a linear path at comfortable speed. Observers thus experienced both natural unencumbered walking and the natural circular relationship between sensory input and motor output that is inherent to our everyday experience.

We hypothesised that visual detection performance would be enhanced in the visual periphery while walking, in accordance with previous research (Cao & Händel, 2019). We additionally hypothesised that oscillations in visual detection performance within the stride-cycle would show distinct phases for targets at separate visual field locations. To preview our results, we found average visual performance was poorer for peripheral compared to central targets, and that stride-cycle based oscillations were reduced in amplitude and phase-lagged for targets at the periphery relative to parafoveal locations.

## Methods

### Participants

Participants were 40 undergraduate psychology students from the University of Sydney who participated in exchange for course credit. Six participants were excluded, as per criteria below, leaving a final sample of *N* = 34. The large sample size was selected due to the lack of prior studies examining modulations within the gait cycle, difficulties in determining an appropriate sample size for power, and to ensure that potential effects would be detected. Each participant provided informed consent before commencing and had normal or corrected-to-normal vision. The study protocol was approved by the University of Sydney Human Research Ethics Committee (HREC 2021/048).

### Apparatus and Materials

An HTC Vive Pro Eye head-mounted display (HMD) was used to display the virtual environment and two wireless HTC Vive Controllers (2018) were used for collecting responses. The HMD contained dual 3.5-inch high-resolution OLED displays (1440 x 1600 pixel resolution, 90 Hz refresh rate) with 110-degree field of view. Positions of the HMD and controllers were tracked in three-dimensional space at 90 Hz resolution, using five HTC Vive Base Station 2.0’s enclosing an open 4.5 m x 12 m rectangular space. To allow the utilization of such a large space and unencumbered walking conditions, the virtual reality environment was transmitted to the HMD using the HTC Vive Wireless Adapter for Vive Pro. The sRanipal Runtime (Version 1.3.2.0; SDK 1.3.6.8) was used to handle eye tracking data collection by the integrated Tobii eye-tracker within the HMD, sampled at 90 Hz.

The virtual reality (VR) environment and experimental procedure were designed and presented within Unity (Version 2020.3.14f1), using the SteamVR Plugin (Version 2.7.3; SDK 1.14.15) on a Windows 11 PC (2 × 16 GB, DDR5 4400 MHz) with a 12^th^ Gen Intel Core i7- 12700K processor (3.60 GHz), and an NVIDIA GeForce RTX 3070 (8 GB, GDDR6) graphics card. The VR environment consisted of a clear space, that matched the dimensions of the tracked physical space, set in an open outdoor scene sparsely populated with trees with a singular simulated natural light source. The trees, ground texture, and skybox used to create the outdoor environment were all free assets available on the Unity Asset store.

All task-relevant visual stimuli were presented on a grey (RGBA 102, 102, 102, 255; 0.4, 0.4, 0.4, 1.0) rectangular screen with simulated dimensions of 13.1° × 50.9° based on a 1 metre viewing distance (see Figure 1b). The target stimulus was an ellipsoid shape randomly orientated 45° clockwise or anticlockwise from vertical subtending 1.2° along its major axis. The QUEST (Watson & Pelli, 1983) adaptive staircase algorithm was implemented to adjust the target contrast (between 0.4, matching the background, and 1.0) that produced an overall accuracy rate of 75% for each stimulus condition. There were four independent staircases for left central, right central, left peripheral and right peripheral targets; Figure 1d displays an example change in contrast at central and peripheral locations. The initial slope (β) was set to 3.5, chance rate (γ) to .5, and lapse rate (δ) to .01 for each staircase, which had resolutions of .004 for the range 0.4 to 1. As the QUEST procedure settled at 75% accuracy after ∼40 targets (after walking ∼5-6 lengths of our 9.5 m path), a jitter was added to each trial’s contrast to enable better estimates of the slopes of the psychometric functions per target condition (see Figure 1e). This jitter was provided by the standard deviation of the prior probability density function of the staircases in real time, resulting in the contrast of the target being selected from the following array of positions: [*M* – 2*SD, M* – *SD, M* – *SD, M, M* + *SD, M* + *SD, M* + 2*SD*] based on the mean (*M*) of the distribution of QUEST-derived threshold estimates in each staircase. The positions one standard deviation from the mean were twice as likely to be sampled to provide an accurate estimate of participants’ 75% detection thresholds (see Figure 1f) and just noticeable difference centred at 75% accuracy (see Figure 1g).

**Figure 1.**
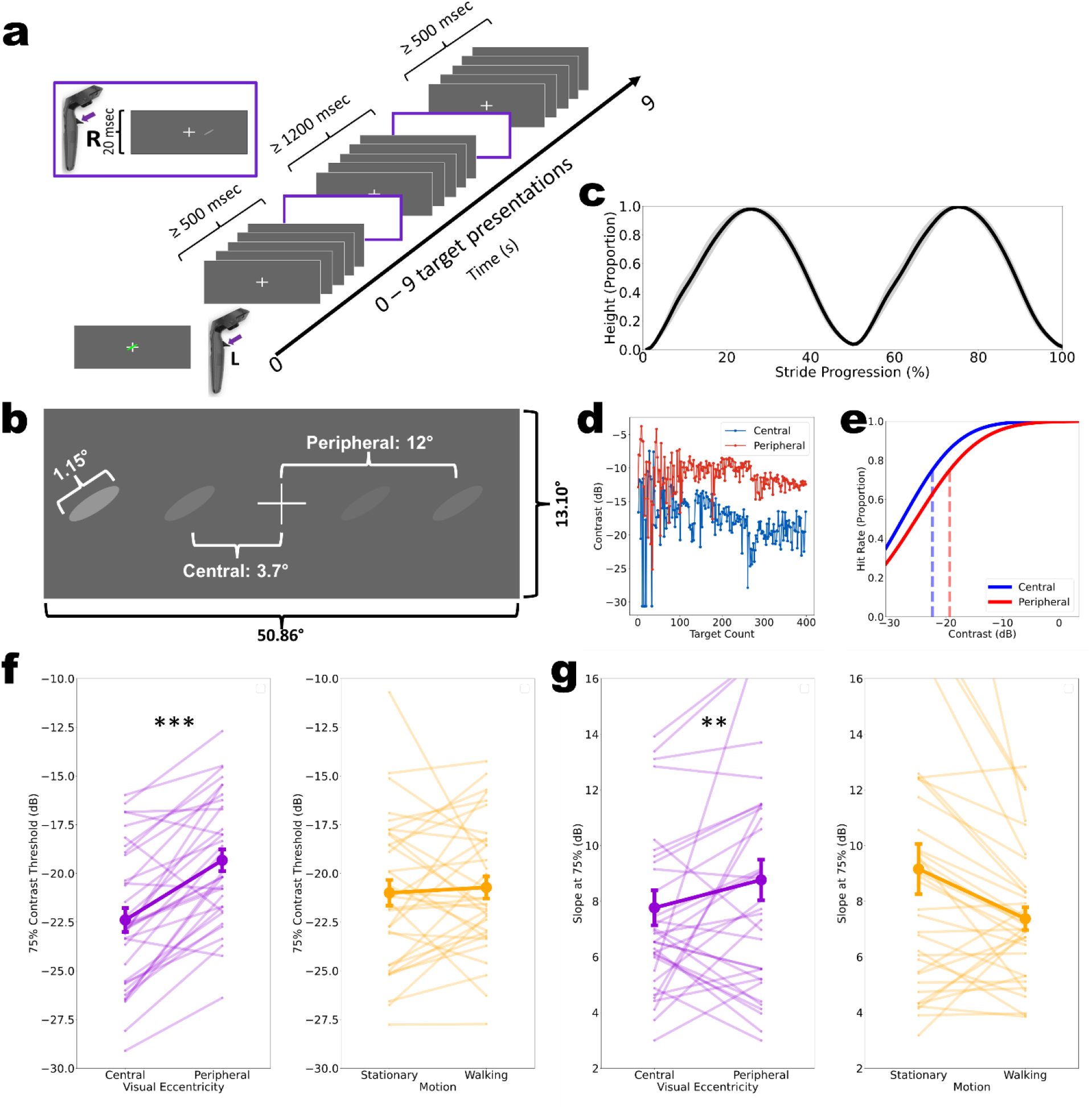
Example of procedure and preliminary data analytics. **a** Example trial sequence while completing a 9 second walking trial. The presentation of a target and response window is outlined in purple. The example sequence has two target presentations, indicated by purple outlined box, with minimum inter-target intervals of 1200 msec. Each walking trial could have between 0 and 9 targets presented, however, no presentations occurred during the first and last 500 msec of each 9 s trial. **b** Schematic of display parameters with the possible target presentation locations (not to scale). Four possible contrasts examples are provided, one per location, yet during the experiment only one target appeared at a time. **c** Each stride, comprised of two steps, were resampled to a normalised range of 1-100% for each participant (*N* = 34). The grand average of the participant head heights was calculated and normalised between the lowest and highest heights to the range of 0 – 1. **d** Example data demonstrating the change in contrast for presentation of central and peripheral targets, in blue and red respectively. The QUEST staircase settled at a lower relative contrast threshold for central targets compared to peripheral targets. **e** Psychometric function fitted to group mean data for central and peripheral targets, in blue and red respectively (see Apparatus and Materials). **f** Group mean (*N* = 34) of 75% contrast thresholds (**α**) derived from fitting psychometric functions to the Quest data for each individual (see Apparatus and Materials). Thresholds were significantly higher for peripheral compared to central locations, *** *p* < .001. **g** Group mean (*N* = 34) of the contrast change required for perceived change (**β**; slope parameter) centred at 75% detection accuracy derived from fitting psychometric functions to individual performance across experienced target contrasts (see Apparatus and Materials). Slopes were steeper when standing still, compared to when walking, ** *p* < .01.

### Procedure

Participants were provided with a participant information statement, followed by an opportunity to ask any questions before they completed the consent form. All participants were informed that their consent and participation could be revoked at any time during the procedure if they wished to not provide their data or did not wish to complete their participation without the need to provide a reason. They were then introduced to the wireless VR apparatus, hand-controllers, power bank, the tracked physical space, and the initial eye tracking calibration procedure before being fitted with the apparatus and given the hand-controllers.

Once the VR program was initiated, participants completed a 4-point calibration process for the HMD’s in-built eye trackers, handled by the eye tracking software. Additional calibration was performed after any potential events that may have affected the accuracy of the tracking (e.g., signal drop-outs, breaks). Participants were then told to situate themselves in front of the virtual display screen, standing on a virtual red cross placed 1 m away from the screen in the centre of the virtual environment, and to read the task instructions. The instructions outlined the flash detection task and the requirements to always fixate on the fixation cross and to make speeded responses via trigger pull on the right-hand controller whenever they detected the target. Participants were also informed that to initiate each 9 second sequence, indicated by a green target on the fixation cross, a left-hand trigger pull was required (Figure 1a). They were then asked to describe the task they were to complete to verify their understanding. If correct in understanding, the participant was instructed to begin the first of three 9 sec sequences while standing still. Otherwise, the participant was assisted in their understanding by alternate verbal instructions until they understood the task correctly and given the opportunity to repeat the practice trial sequence. During this practice the targets presented were easy to detect, with a fixed supra-threshold contrast (adaptive staircases were not updated).

Before beginning their first block of walking sequences, participants were instructed to stand on a red cross repositioned to indicate the starting location of the walking sequence. The task was identical whilst walking, maintaining central fixation and with the additional instruction to maintain approximately 1 m between themselves and the display as it moved along a straight line at constant speed set by the walking guide.

Walking speed was set by the virtual walking guide, an animated 3D object in the virtual environment positioned in front of the observer which traversed a 9.5 m distance in 9 seconds and served as a pacesetter for the participant. Following the walking guide resulted in a walking speed of 1.06 m/s. Although slightly slower than typical human walking speed of approximately 1.4 m/s over long distances in natural environments (Hausdorff et al., 1996; Matthis et al., 2018), this reduced speed was found to be appropriate and comfortable based on the unfamiliarity some participants had with walking freely in a VR environment (Davidson et al., 2024). During each 9 second walking trial, targets were displayed to the left or right of fixation by 3.7° in the central condition (with no vertical offset) and by 12° in the peripheral condition (see Figure 1b). All locations were jittered by 0.5° horizontally. Crucially, simultaneous eye-movement recordings enabled the eccentricity of each target relative to fixation to be calculated, with subsequent exclusion from analysis when targets were not presented within our predefined central and peripheral regions of interest. Participants’ detection accuracy and responses were collected as measures of visual detection ability and were analysed as outlined below (see Figure 1). The time between target onset and response was taken as response time and the number of responses at each time-point across the stride-cycle taken to reflect a response likelihood.

During sequences in walking blocks, the virtual display screen would smoothly traverse linearly along the walking path at a constant velocity and height, beginning motion when the participant started the sequence (i.e., left-hand trigger pull). For the first two sequences of the block, the participant was accompanied to ensure their safety, and to verify this approximate distance was maintained comfortably whilst walking. Upon traversing the walking path, the participant was required to turn 180° before starting the following sequence, such that they returned on the path they had walked. Once comfortable with the procedure while walking, the participant was allowed to complete the remainder of the experiment independently, initiating the sequences themselves with left-hand trigger pulls.

Participants completed 10 blocks, five stationary blocks and five walking blocks, with the order of the blocks randomised except the first block. Each block had twenty 9-s sequences, each of which had 7, 8, or 9 targets presented relative to predefined times, with each target having a 10% chance of being withheld from presentation. Targets were presented in one of four possible locations as per the factorial design, either left or right of fixation and either centrally (± 3.7°) or peripherally (± 12°), with the inclusion of a ± 0.5° horizontal jitter. Targets were presented for 20 ms and were spaced by variable intertrial intervals with a minimum of 1200 ms. Targets were never presented during the first and last 500 ms of sequences.

### Data Analyses

Each sequence provided 3D time-series data (x, y, z coordinates) for head position, target position, gaze origin, and gaze direction. Individual steps were extracted based on head height, which follows a roughly sinusoidal pattern during walking. Peaks and troughs in vertical head position correspond to the approximate swing and stance phases of the stride-cycle, which we identified using a peak detection algorithm. Changes in head height during all strides (two sequential steps) were averaged for all participants, as displayed in Figure 1c, and then normalised between the lowest and highest value to a range of 0-1. Target performance data (i.e., hits, misses, false alarms and response times) were mapped according to target onset relative to the nearest percentile of the stride-cycle. All analyses were performed using custom Python (version 3.11.1) code, and ANOVAs were performed in JASP (version 0.18.0.0).

### Psychometric Function Fitting

Psychometric functions were fitted to the participant data using a cumulative normal distribution with maximum likelihood estimation. From each function, the 75% threshold (α) and slope (β) parameter were retained. The threshold values are reflective of the participants’ sensitivity to visual stimuli with greater values indicating greater difficulty (see Figure 1e).

### Eye Movement Data

At each time point gaze coordinates were calculated using gaze origin and direction values, facilitating the calculation of target eccentricity. Individuals’ eye gaze data were projected onto the simulated display screen as polar coordinates and used to verify correct fixation.

Specifically, the proportion of frames where an individual deviated from the fixation cross (exceeding 1° eccentricity) during each 9 second sequence was used to quantify overall fixation. One participant was removed for recording a 75% proportion of deviation from fixation. Using the same projection procedure, the eccentricity of the targets relative to fixation at the moment of stimulus onset was also calculated. We retained central targets for analysis when they were presented within 7° of eccentricity from gaze fixation, and peripheral targets for analysis when presented between 7 and 20° of eccentricity from fixation. Any targets that were presented within 0.5° or beyond 20° of eccentricity from the locus of fixation were excluded as potential blinks or extraneous eye-movements.

### Gait Extraction

Stride-onsets were extracted based on the time-series of the vertical head position as described in (Davidson et al., 2024). As walking shifts the centre of mass sinusoidally (Hirasaki et al., 1999; MacNeilage & Glasauer, 2017), troughs on the vertical axis of motion correspond to the double support stance phase of the gait cycle. Stride-lengths were normalised for analysis by resampling the time-series data to 100 points (taken as percentage of stride-cycle progression; see Figure 1c). Our categorisation of stance and swing phases was approximated as per the findings of MacNeilage and Glasauer (2017), attributing the ∼1-10%, ∼40-60%, and ∼90-100% of the gait progression to the stance phase. These swing and stance phases are displayed with grey shading in our visualization of stride-cycle based effects (see Figure 2 and Figure 5).

**Figure 2.**
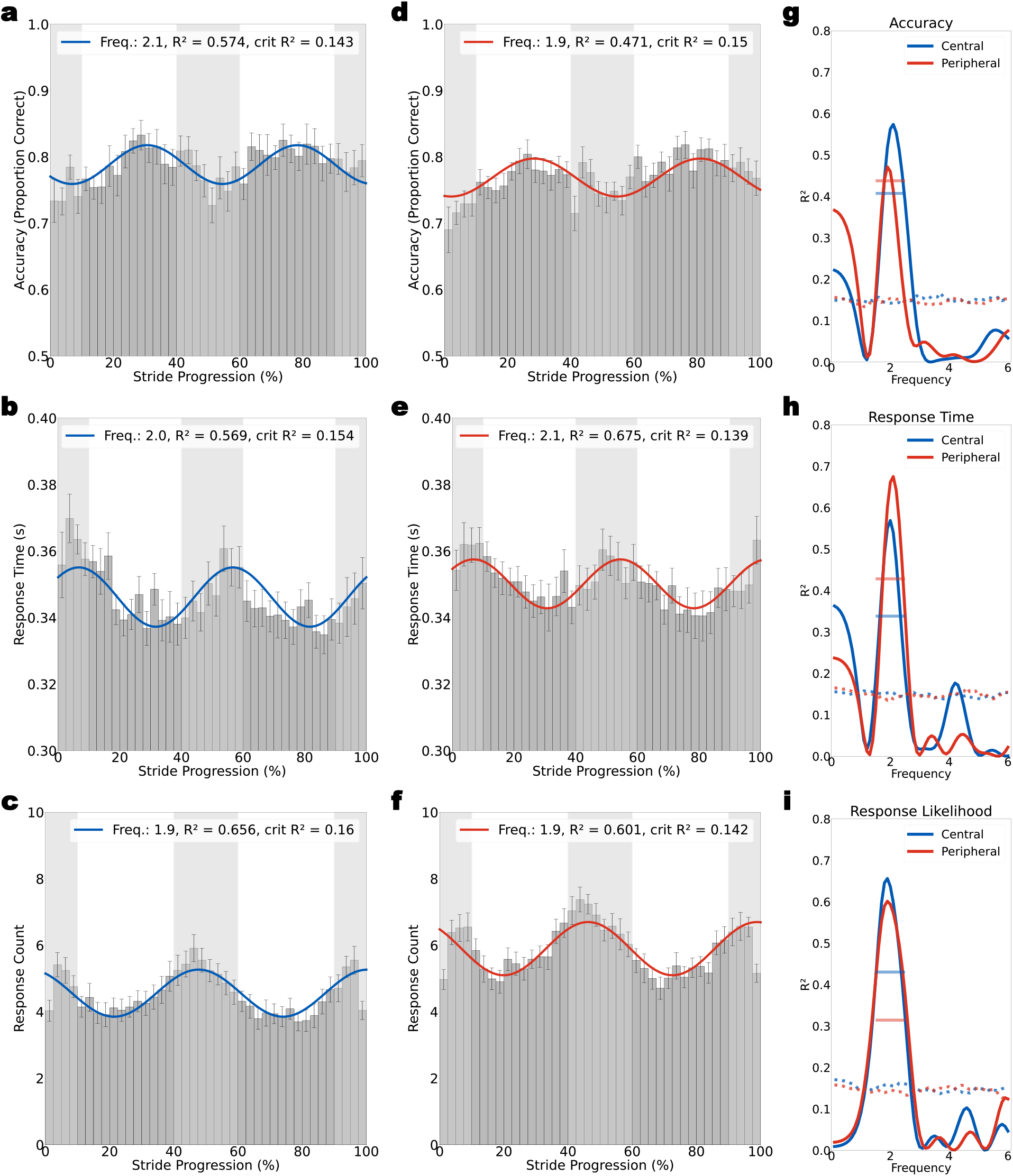
Oscillations in performance at central and peripheral target locations within the stride-cycle. **a, d** Group-level (*N = 34*) accuracy data show oscillations at 2.1 cps and 1.9 cps for centrally and peripherally presented targets, respectively. Best-fitting Fourier models are displayed as coloured lines, blue in left columns for centrally presented targets and red for peripherally presented targets. Light and dark grey regions indicate the estimated stance and swing phases of the stride-cycle, respectively. **b, e** Group-level reaction time data show oscillations at 2.0 cps and 2.1 cps for centrally and peripherally presented targets, respectively. **c, f** Group-level oscillations in response likelihood show oscillations at 1.9 cps. **g-i** Fourier models were fit at a fixed frequency between 0.1-6 cps (in steps of 0.1) on the group-level data (*N = 34*). The solid blue and red lines display the goodness-of-fit (*R*^2^) calculated for each performance measure at central and peripheral target locations, respectively. Dotted lines show the upper 95^th^ percentile of *R*^2^ values at each fitted frequency obtained from a null distribution of group-level data shuffled in time (*n* = 1000 permutations). Solid horizontal lines show the upper 95^th^ percentile when retaining the maximum *R*^2^ value across all frequencies in the range 1.5- 2.5 cps per permutation (a more conservative test, guided by the results and frequency range of interest in Davidson et al., 2024)

### Performance Relative to Stride

All target onsets were allocated to the percentile (from 1-100%) of the stride they occurred in. For stride analyses, performance was averaged for targets occurring within 40 linearly spaced bins of 2.5% width, with zero overlap. We analysed detection accuracy, taken as the proportion of correct detections (i.e., hits) out of total presented targets, and response times relative to target onset within each stride-cycle. We additionally analysed the likelihood of committing a response (trigger pull) as a function of stride-cycle phase.

We tested for significant oscillations in each measure at both the participant and group level. For each measure, a Fourier series model was fitted using the equation:

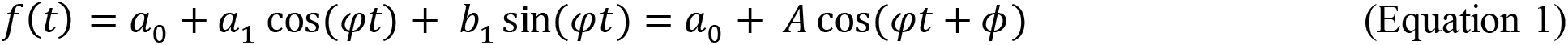

where ***φ*** is the periodicity (cycles per stride), ***t*** is the progression of the stride (as a percentage), ***a*_1_** and ***b*_2_** are the coefficients of the cosine and sine components and ***a*_0_** is a constant, the central value of the oscillation. The resulting fits had amplitudes of ***A*** and shifts in phase of ***ϕ***. At the group level, we retained the Fourier model with maximum goodness of fit when stepping from .1 to 6 cps, in increments of .1 cps (displayed as an overlay in Figure 2). For tests of statistical significance, the goodness of fit (*R*^2^) at each frequency (cps) was used as an indication of oscillation strength. For statistical purposes, a null distribution of *R*^2^ values was calculated by first shuffling the labels for each percentile bin (from 1-100% with replacement), before repeating our main analysis. Specifically, on each permutation (*N* = 1000), we repeated our binning procedure and refit the Fourier series model over the entire frequency range of interest (.2 to 10 cps, in .2 steps). The 95^th^ percentile of *R*^2^ values at each frequency (see Figure 3, dotted lines) was used to test for significance by comparing the *R*^2^ of the original data to the 95^th^ percentile of the permuted data, with significant oscillations at a given frequency indicated when the original *R*^2^ exceeds the 95^th^ percentile of the permuted data.

**Figure 3.**
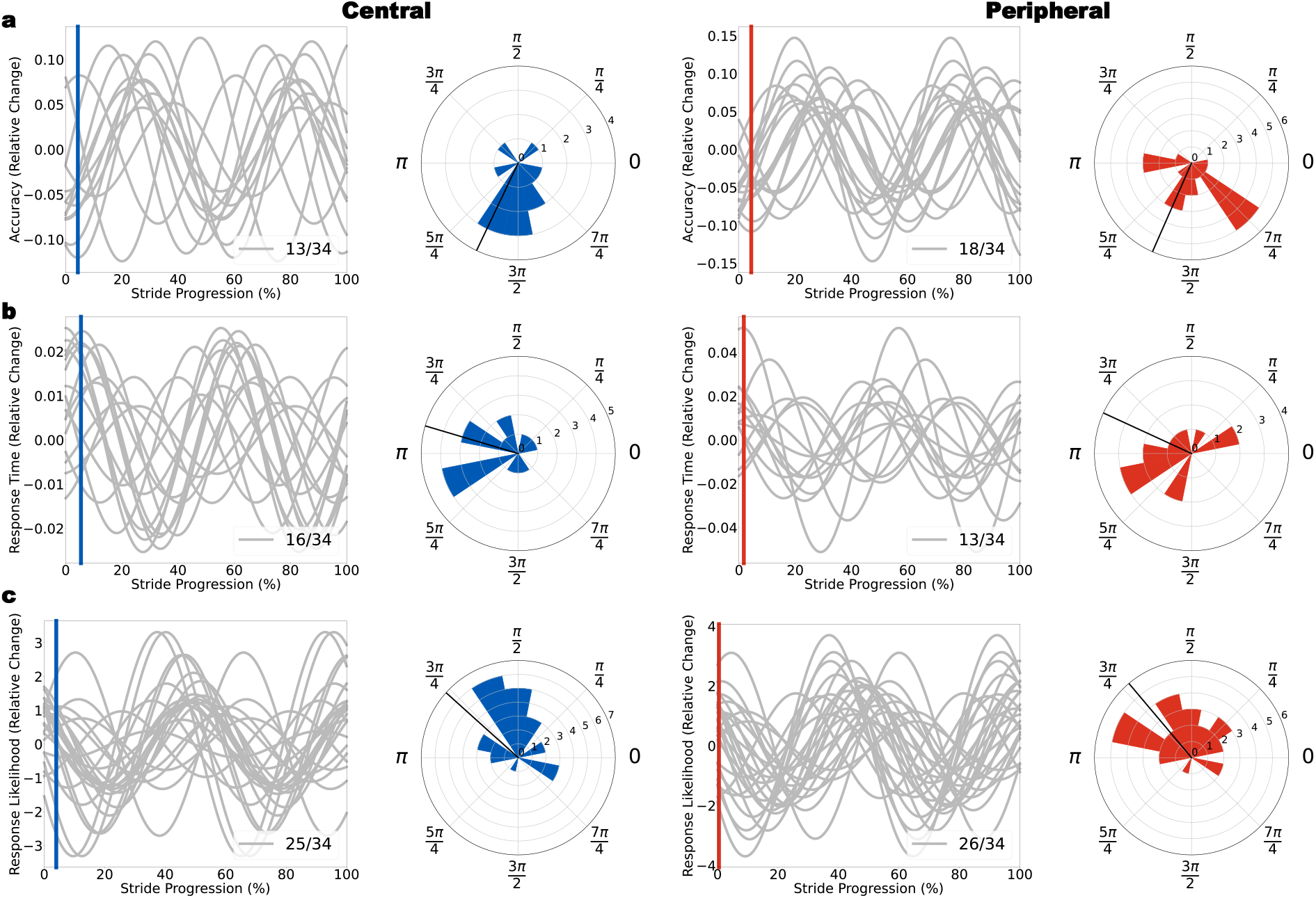
Oscillations in Participant-level Performance. *Note.* Oscillations for centrally presented targets are presented in blue (left panels) and oscillations for peripherally presented targets are presented in red (right panels). Black radial lines on polar plots represent the mean phase for the subset of participants with significant oscillations. For each performance measure, the modal frequency was fitted for all participants. **a** *n* = 13 participants displayed significant oscillations in accuracy (1.5–2.5 cps) for centrally presented targets. The fit at modal 1.8 cps is displayed, with phase clustering shown in the polar plot. Right column shows accuracy oscillations following peripheral targets (*n* = 18). **b** Displays the subset of participants with significant oscillations in response time for central (*n* = 16) and peripheral targets (*n* = 13) fit at the modal frequency of 1.8 cps. **c** Displays participant oscillations in response likelihood following central (*n* = 25) and peripheral (*n* = 26) targets.

### Jackknife analyses to compare central and peripheral oscillations in performance

We observed strong oscillations in central and peripheral performance at the group-level (see Figure 2) and recognised that these were being driven by a subset of participants in our sample (see Figure 3). We were particularly motivated to statistically quantify whether a reliable phase-shift was present between central and peripheral locations, as previous research has shown similar examples of rhythmically induced perceptual oscillations (Fakche & Dugué, 2024; Lozano-Soldevilla & VanRullen, 2019; Sokoliuk & VanRullen, 2016). To compare the parameters of our Fourier fits on performance at central and peripheral locations, we applied the same fitting procedure as described above using a jackknife resampling procedure and super subject resampling procedure. Specifically, the best-fitting group-level Fourier fits were obtained for each permutation using *N*-1 subsets of our total sample. For accuracy, reaction times, and response likelihoods, we retained the phase and amplitude parameters of our jackknifed distributions for statistical comparisons. The results of this procedure are displayed in Figure 4.

**Figure 4.**
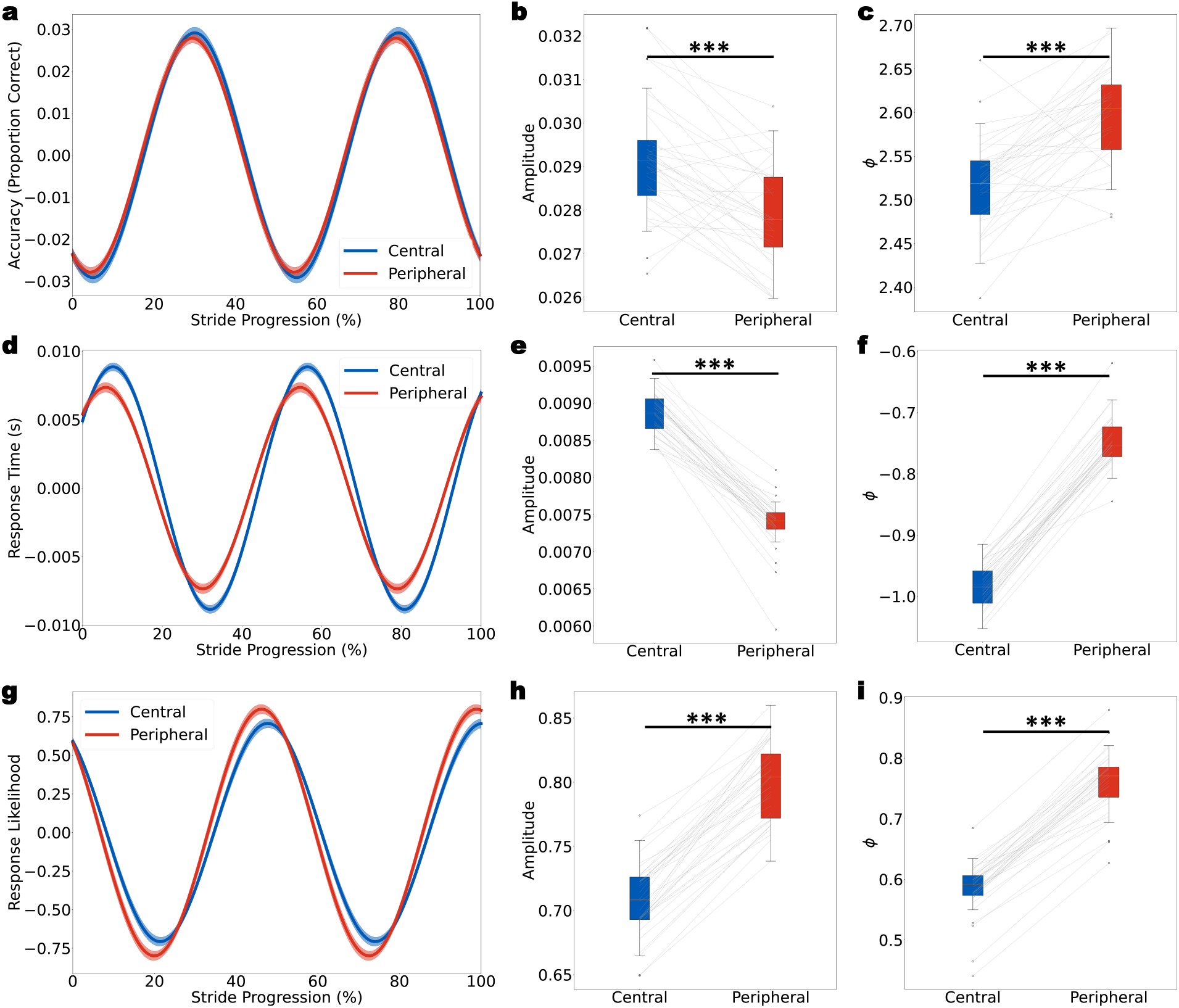
Oscillations are phase lagged at peripheral compared to central locations. **a** Average Fourier models for oscillations in accuracy at central (blue) and peripheral locations (red) obtained via jackknifed (*N*-1) distributions. Shading represents ±1 SEM. **b-c** Distribution of amplitude estimates for oscillations in accuracy (b) and phase (c) at central (blue) and peripheral (red) target locations. Oscillations were smaller in amplitude, and earlier in phase at central locations. Box and whisker plots represent the distribution of the estimates with the interquartile range indicated by the whiskers. **d-f** Similar oscillations with larger amplitudes and earlier phases for reaction time data at central locations in comparison to peripheral locations. **g-i** Responses to peripheral targets oscillated at a higher amplitude, and an earlier phase than central target responses. *** *p* < .001

**Figure 5.**
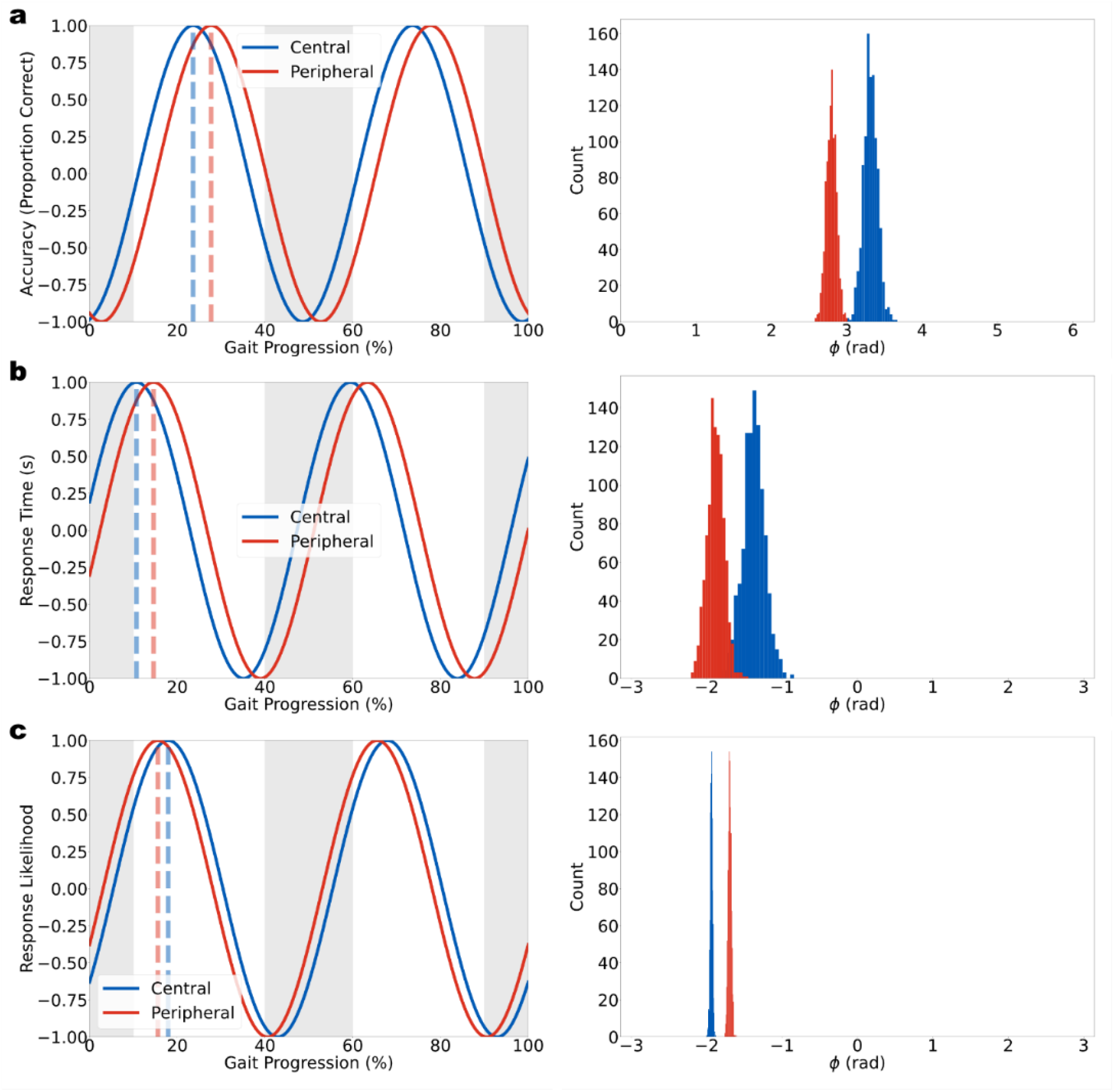
Normalised Oscillations of Super Subject. *Note.* Plot of normalised oscillations of a super subject comprised of all participants with significant oscillations for both centrally and peripherally presented targets for each performance measure. Dashed lines indicate the peak of oscillations. Frequency distributions of the phase shift (ϕ) for 1000 bootstrapped resamples of the super subject dataset are displayed. **a** Oscillations in accuracy of super subject, fitted at 2.0 cps. **b** Oscillations in response time of super subject, fitted at 2.05 cps. **c** Oscillations in response likelihood of super subject, fitted at 1.9 cps.

### Exclusion Criteria

Participants with data from less than four walking or stationary blocks were excluded for insufficient data collection (*n* = 1). Head position data was visually inspected for aberrant sequences and abnormal strides that may have occurred due to hardware malfunction, signal drop-out, or abnormal walking patterns (e.g., limping). Across all participants, an average of 0.57 sequences (*SD* = 0.85, range 0 – 4) and an average of 0.43 steps (*SD* = 1.88, range 0 – 11) were excluded using this procedure. No participants were excluded based on this procedure.

Participants determined not to have fixated (i.e., kept gaze within 1°), according to the data analysis process, for at least 75% of their collective data points were excluded (*n* = 1).

Additionally, those with overall accuracy for each adaptive staircase less than 65% (*n* = 3) or greater than 85% (*n* = 1) were excluded for possible floor and ceiling effects, respectively.

Specific targets were excluded from analyses if they were closer than 0.5° (*M* = 5.58 targets, *SD* = 5.17, range 0 – 23) or exceeded 20° in visual eccentricity (*M* = 120 targets, *SD* = 217, range 0 – 982), most being too slow in walking or not fixating properly, respectively.

## Results

### Visual detection psychometric functions interact with motion type and target location

We designed a visual detection task to compare performance at central and peripheral visual field locations while walking in a wireless VR environment. A 2 (Motion: Stationary vs Walking) x 2 (Visual Eccentricity: Central vs Peripheral) repeated-measures ANOVA compared the contrast thresholds (α at 75% detection; see Figure 1e) and slope parameters (β; see Figure 1f) derived from individual psychometric functions. There was no main effect of motion on participants’ detection thresholds when comparing standing still to walking (*F*(1, 33) = 0.25, *p* = .62, η^2^ = .002). There was a main effect of eccentricity, such that participants’ detection thresholds were lower for centrally presented targets than for peripherally presented targets (*F*(1, 33) = 53.96, *p* < .001, η^2^ = .280). The interaction between motion and eccentricity was not significant (*p’s* > .05).

When comparing the effects of motion and visual eccentricity on the slope parameter of psychometric functions (see Figure 1f), there was a main effect of motion, such that slopes were steeper when standing still than when walking (*F*(1, 33) = 7.67, *p* = .009, η^2^ = .06. There was no main effect of eccentricity on the slope parameter (*F*(1, 33) = 3.25, *p* = .08, η^2^ = .02), and no interaction (*p* > .05).

An additional three-way 2 (Visual Eccentricity: Central vs Peripheral) x 2 (Visual Field: Left vs Right) x 2 (Step Foot: Left vs Right) repeated-measures ANOVA was performed on the accuracy and response times from target presentations during walking. There were no significant main effects nor any significant interactions for either accuracy or response times (see Table 1), *ps* > .05.

**Table 1.** Mean Accuracy and Response Times by Experimental Condition.

| Visual Eccentricity | Visual Field | Step Foot | Accuracy |  | Response Time |  |
| --- | --- | --- | --- | --- | --- | --- |
|  |  |  | *M* | *SD* | *M* | *SD* |
| Central | Left | Left | .78 | .11 | 348 | 29 |
|  |  | Right | .77 | .09 | 352 | 35 |
|  | Right | Left | .78 | .10 | 347 | 26 |
|  |  | Right | .78 | .07 | 352 | 22 |
| Peripheral | Left | Left | .77 | .06 | 355 | 20 |
|  |  | Right | .77 | .07 | 354 | 22 |
|  | Right | Left | .76 | .06 | 348 | 25 |
|  |  | Right | .76 | .05 | 350 | 26 |

*Note.* The proportion of correct responses to target presentations was taken as a measure of accuracy. The response time is presented in milliseconds (ms).

Together, these results demonstrate overall that thresholds, accuracy and reaction times were matched in our conditions, owing to the successful application of our QUEST staircase to equate performance. Notably however, there was an effect of target location on the contrast thresholds reached to match performance. Contrast thresholds decreased in the centre relative to the periphery to maintain 75% detection.

### Stride-cycle phase modulates performance for targets at central and peripheral locations

In addition to comparing average performance between central and peripheral target locations while walking, our wireless VR environment enabled an analysis of changes in performance relative to the phases of the stride-cycle. When binning target-onsets to their relative stride-cycle phase (see Methods), accuracy and reaction times at both central and peripheral locations showed significant oscillations at ∼2 cps at the group-level (see Figure 2). Response likelihood was also modulated at ∼2 cps. Oscillations in accuracy occurred such that participants had improved detection for targets presented whilst they were in the swing phase of their steps (see Figure 2a, d). Similarly, targets presented during the swing phase were also responded to faster than those in the stance phase (see Figure 2b, e). When aligning response onsets to stride-cycle phase, response likelihood was greatest in the early stance phase, and lowest in the swing phase (see Figure 2c, f).

We tested the statistical significance of these group level oscillations using a two-step permutation procedure (see Methods). In brief, the observed fit-strength at a given oscillation (e.g., *R*^2^ value at 2 cps) was compared to the upper 95^th^ percentile of *R*^2^ values obtained from fitting the same Fourier model to permuted data (*n* = 1000 permutations). As shown in Figures 2 g-i, the *R*^2^ values from the Fourier models fit to observed data at ∼2 cps far exceeded the upper 95^th^ percentile of *R*^2^ values expected by chance. This indicates the highly oscillatory nature of the participants’ performance over the gait cycle.

### Participant level analysis of oscillations at central and peripheral locations

We next performed participant-level analyses of oscillations according to stride-cycle phase to determine the prevalence and phase-consistency of oscillations at central and peripheral target locations. For this, only participants with significant oscillations in the range 1.5–2.5 cps were included, and to estimate the phase-clustering, each participant was fit with a Fourier model with a fixed frequency set to the mode of significant oscillations within each subset of significant participants. For oscillations in accuracy, to central (*n* = 13) and peripheral targets (*n* = 18) the mode was 1.8 cps. The distribution of phase angles at both locations was significantly non-uniform (Rayleigh’s test of non-uniformity; central *Z* = 4.07, *p* = .014; peripheral *Z* = 5.57, *p* = .002). Figure 3a displays the individual sinusoidal fits to accuracy at the mode of 1.8 cps for both central and peripheral locations, as well as polar plots of phase consistency.

The oscillations in response time (central *n* = 16, peripheral *n* = 13) had the same modal frequency (1.8 cps). However, these phase distributions were not significantly non-uniform (central: *Z* = 2.64, *p* = .070; peripheral: *Z* = 2.09, *p* = .123, see Figure 3b). A large portion of our sample demonstrated significant oscillations in response likelihood (central *n* = 25, peripheral *n* = 26). The distribution of these phases were significantly non-uniform following targets at both locations (central, *Z* = 7.21, *p* < .001; peripheral, *Z* = 6.96, *p* < .001; see Figure 3c).

### Oscillations in performance are reduced in amplitude and phase-lagged at peripheral compared to central locations

Finally, we tested whether oscillations in performance at central and peripheral locations differed in either amplitude or phase using a jackknife resampling procedure (see Methods). This method repeated the group-level Fourier fitting as described above on each permutation of *N*-1 subsets of our total data. For accuracy, reaction times, and response likelihoods, we created a distribution of amplitude and phase parameters from this permutation and tested for consistent differences between locations using paired-samples *t*-tests. All jackknife permutations were fitted with the mean frequency between locations of the group-level Fourier fits (i.e., 2.0 cps for accuracy, 2.05 cps for response time, and 1.9 cps for response likelihood).

Figure 4 displays the mean group-level Fourier fit obtained from all *N*-1 jackknifed estimates. Oscillations in accuracy following central targets were significantly greater in amplitude than oscillations to peripheral targets (*t*(33) = 5.07, *p* < .001, *d* = 0.87). Oscillations in central accuracy were phase-lagged relative to the peripheral targets (*t*(33) = −6.58, *p* < .001, *d* = −1.13). The same pattern was present for the reaction time data, with significantly greater oscillation amplitude at centrally presented targets compared to the periphery (*t*(33) = 21.41, *p* < .001, *d* = 3.67), which were also earlier in phase than peripheral target locations (*t*(33) = - 33.26, *p* < .001, *d* = −5.70). When examining differences in response likelihood following central or peripheral targets, oscillations to peripheral targets were significantly larger in amplitude (*t*(33) = 22.02, *p* < .001, *d* = 3.78), and also delayed in phase (*t*(33) = −29.98, *p* < .001, *d* = −5.14).

### Super subject analysis for within-participant comparisons

We additionally performed a complementary super subject analysis to confirm these differences in oscillatory features between central and peripheral locations. This analysis focused only on the subset of participants which had significant oscillations at both central and peripheral locations (accuracy *n* = 6, response time *n* = 6, response making *n* = 21). Importantly, by including only those participants, this analysis mitigates the risk that the differences we report at each eccentricity may be driven by comparisons between unique subsets of participants (such as those with significant oscillations at only one eccentricity).

To test whether there were differences between the phases (ϕ) of central and peripheral performance oscillation models, a super subject dataset was made for each performance measure composed of participants who showed significant oscillations at ∼2.0 cps (1.5-2.5 cps) at both eccentricities. For example, the super subject datasets contained all target presentations for the six participants with oscillations at central (*n* = 3374 trials), and peripheral locations (*n* = 4518) trials). The super subject performance was fitted for oscillations at the mean of the best fitting frequencies for both eccentricities of the respective group-level performance oscillations (accuracy *φ* = 2.0, response time *φ* = 2.05, response making *φ* = 1.9; see Figure 5). The super subject datasets were then resampled via bootstrap (with replacement) 1000 times, excluding a portion equivalent to a single participant on each iteration (accuracy prop. = 1/6, *n* = 1315 trials; response time prop. = 1/6, *n* = 983 trials; response making prop. = 1/21, *n* = 1031 trials). Each bootstrapped dataset was fitted at the same frequency as their respective super subject dataset and the distribution of their phases (see Figure 5) compared to test for phase differences between oscillations at the two eccentricities. Two-sample Kolmogorov-Smirnov tests were performed on the distribution functions. The phases for the central accuracy oscillations (mode = 3.31 radians) were distributed significantly higher than those for the peripheral accuracy oscillations (mode = 2.80 rad), such that accuracy peaked 8.19% earlier in the gait cycle for central targets, *D* = .999, *p* < .001 (see Figure 5a). A similar difference in phases of the central (mode = −1.37 rad) and peripheral (mode = −1.88 rad) response time oscillations was found, where response times peaked 8.14% earlier in the gait cycle for central targets, *D* = .956, *p* < .001 (see Figure 5b).

However, the distribution of phases for peripheral response likelihood oscillations (mode = −1.94 rad) were instead significantly higher than those for the central response likelihood oscillations (mode = −1.69 rad), such that the likelihood of a response being made peaked 3.85% earlier in the gait cycle for peripheral targets, *D* = 1.000, *p* < .001 (see Figure 5c).

Together, these within-participant comparisons confirm the results of our group-level jackknife procedure. Specifically, oscillations in visual detection accuracy and reaction time are larger in amplitude and earlier in phase for targets presented at central compared to peripheral target locations.

## Discussion

The current study examined the effects of walking on visual detection performance, with a specific focus on changes in performance at central and peripheral visual field locations within the stride-cycle. Participants made speeded trigger-pull responses with their right hand to indicate their detection of a presented target. When comparing overall performance, we observed an interaction between the threshold and slope parameters of psychometric functions based on whether targets were in the centre or periphery, with increased detection thresholds for peripheral targets. Pronounced oscillations in task performance were also entrained to the rhythm of the stride-cycle, with a significant difference in the amplitude and phase lag of oscillations at central compared to peripheral target locations.

### Performance oscillations at central and peripheral locations are entrained to the stride-cycle

Contrast thresholds increased at peripheral compared to central locations, supporting prior observations of higher contrast thresholds for peripheral visual targets (Cannon, 1985; Rijsdijk et al., 1980; Rovamo & Virsu, 1979; Virsu & Rovamo, 1979). When examining changes in performance relative to stride-cycle phase, strong oscillations were present in accuracy and reaction time for both central and peripheral targets. These results confirm the modulatory effects of walking on visual perception observed in other studies (Cao & Handel, 2019; Chen et al., 2022a; 2022b) and adds to recent work showing more specifically that perceptual modulations are linked to the stride-cycle (Davidson et al., 2023; 2024). Consistent with Davidson et al. (2024), within-stride oscillations were found for all three performance measures: accuracy, response time, and response likelihood. These group-level oscillations were all at a frequency of ∼2 cps with improved accuracy and reaction times in the swing phase of the stride-cycle, and responses most likely around the time of heel-strike.

### Participant level oscillations cluster but can vary in phase

At the participant-level, we observed significant phase clustering of these oscillations, such that accuracy in most observers improved during the swing-phase of each stride and reaction times were faster. Here, as in Davidson et al. (2024), we note that individual participants who exhibited significant oscillations showed some variability in the precise location of their performance peak. Peaks tended to cluster around the middle of the swing-phase but a small group of outliers showed a counter-phase relationship to the average group-level result and thus showed a performance decrease in the swing-phase. While the group level result is clear, here and in our other study of gait-related perceptual modulations (Davidson, et al., 2024), the reason behind the outlying participants is not clear. Two possible reasons are that those participants who showed clear anti-phase results adopted a different response strategy, or they may have had unusual or idiosyncratic gait patterns. A simple visual inspection of gait patterns (based on head position data) indicated that participants showing anti-phase performance did not differ markedly from other participants. A full gait analysis quantifying spatiotemporal factors and kinematics would provide more insight into this possibility. More studies will be needed to understand the anti-phase performance shown by a minority of observers.

One potential explanation of anti-phase performance that can be excluded is an account based on eye movements towards the targets. In the present study there were only occasional instances of large eye-movements. These will inevitably occur in designs involving peripherally presented targets as participants are sometimes tempted to break central fixation and look directly at the peripheral target. Our participants were instructed not to break fixation and told that the strategy cannot be effective as target duration was just 20 ms and saccades take far longer than this to program and execute (saccadic latencies range from 150 to 200 ms: Smit et al., 1987). Nevertheless, some large saccades still occurred and we applied stringent exclusion criteria to identify and removed the affected trials from the data (see Methods). We can be confident, therefore, that all data included in our analyses involved central fixation. As a result, we can rule out an explanation of these phase differences across participants in terms of differences in eye-movement activity, but other options remain, and future studies will be needed to explore them.

One alternative explanatory mechanism for the variability in participant-level oscillations relates to the demands of locomotion itself. At the beginning (i.e., toe off) and end of a step (i.e., heel strike), the requirement for motor planning and execution shifts sensory focus from the visual modality to comparing motor efference copies with afferent proprioception (Chagnaud et al., 2012; Cullen, 2004; Wolpert & Flanagan, 2001). At these critical stages of the stride-cycle, the unpredictability of head movements is also greatest, potentially shifting requirements toward the vestibular modality for increased proprioception, balance and stability (Bent et al., 2005). It is currently unclear whether differences in baseline balance, proprioception, or dual-task abilities may account for some of the differences in participant level-effects we have observed. A potential manipulation that could serve to investigate this effect would be to heighten the motor requirements of specific portions of the stride-cycle, for example by having participants walk up an inclined path or on an unstable surface. The use of a passive control condition – such as by replaying the locomotion induced optic flow in a motion simulator, could also serve to decouple the motor and proprioceptive demands experienced while walking, to test their respective influence on the oscillations we have observed.

### Performance oscillations differ in amplitude and phase based on visual field location

Both central and peripheral vision exhibited oscillations for each performance measure (accuracy, reaction time and response likelihood), however, there were differences between the amplitudes and phases of the oscillations across the eccentricities. Specifically, when comparing the amplitude and phase of oscillations at each target location, we revealed that oscillations in accuracy and reaction time were significantly larger in amplitude, and earlier in phase, for central compared to peripheral targets. What might drive these smaller and lagged performance modulations for peripheral targets? Prior research has indicated that differences in visual ability across different visual locations are fundamentally tied to the non-uniform distribution of receptors on the retina and primary visual areas (Curcio & Allen, 1990; Rovamo & Virsu, 1979; Virsu & Rovamo, 1979; see Himmelberg et al., 2023 for review). At present however, it is unknown how these asymmetries may map from the tradition of seated experimental settings to the active observer, or whether different regions may be activated asymmetrically during the stride-cycle (Parker et al., 2020).

One possibility advocated by Cao and Händel (2019) is that a trade-off may occur between central and peripheral locations during walking, with the periphery prioritised for safe navigation because potential moving hazards are most likely to enter from the periphery.

Detecting these efficiently would avoid potential collisions and keep the individual reactive to the inevitable changes occurring during locomotion in a dynamic perceptual environment. In our data set, central and peripheral targets were never presented simultaneously, which precludes a direct test of this proposed trade-off, but it remains a tantalising proposition to test in future research. Notably however, oscillations in performance were lower in amplitude for targets at peripheral locations during walking, potentially indicating a minor advantage with respect to variation within the stride-cycle. Future research could test whether these amplitude decrements scale with increasing eccentricity, and at other polar angles from fixation (i.e. off the horizontal axis), to confirm whether a reduction in oscillation depth varies with eccentricity. For example, future work could investigate whether this difference applies to different visual location comparisons, such as vertical (as opposed to lateral) eccentricity (Abrams et al., 2012; Corbett & Carrasco, 2011), perceived depth (Nasr & Tootell, 2018), and between nasal and temporal fields (Paradiso & Carney, 1988; Rafal et al., 1991).

Similarly, recent work has identified that induced perceptual oscillations propagate across the visual field, potentially supported by travelling waves across the cortex (Fakche & Dugué, 2024; Sokoliuk & VanRullen, 2016). Both theta (3-7 Hz) and alpha (8-12 Hz) oscillations have been implicated as modulators of visual and auditory sensitivity (Di Gregorio et al., 2022; Ergenoglu et al., 2004; Fuentemilla et al., 2008; Köhler et al., 2021; Sakowitz et al., 2000; Weisz et al., 2011; Zoefel & VanRullen, 2017) and in divided and focused visual attention (Fiebelkorn et al., 2018; Landau et al., 2018). While we did not explicitly manipulate attention, previous manipulations, such as via a valid or invalid cue, have successfully altered the phase of oscillatory performance on visual detection tasks (Busch & VanRullen, 2010; Fiebelkorn et al., 2018; Ho et al., 2017; Landau & Fries, 2012; Zhang et al., 2019). To understand the extent to which the phase differences we have observed also rely on attentional processes, the incorporation of a cue to central and peripheral locations could be an informative condition to include. Similarly, whether the phase shifts we have observed continue to scale with increasing eccentricity will be an exciting prospect for future research.

## Conclusion

The current study has confirmed the modulatory effect of locomotion upon visual detection, measuring oscillations in visual detection performance at both central and peripheral target locations when controlling for eye-movements. Oscillations in central vision were larger in amplitude, and oscillations in peripheral vision were slightly phase-lagged, suggesting an interaction between the entrainment of performance and visual field locations. Future research may focus on the inclusion of increased target locations in order to characterise whether the differences in phase scale with visual eccentricity. In addition, as modern locomotion regularly incorporates dual-task requirements that may divide attention (such as while holding a phone), an important step will be to determine whether performance oscillations while walking can be biased by an attentional cue.

